# Early Diagnosis of Parkinson via Transient Beta Frequencies in a Delayed Van der Pol Model

**DOI:** 10.1101/2025.09.04.673844

**Authors:** M. A. Elfouly, T. S. Amer

**Affiliations:** Independent Researcher (No Institutional Affiliation) –Egypt; Mathematics Department, Faculty of Science, Tanta University, Tanta 31527, Egypt

**Keywords:** Parkinson’s disease, Transient Beta Bursts, Van der Pol Oscillator, Delay Differential Equations, Early Diagnosis, Neurodynamics

## Abstract

Biological motor circuits are shaped by pathway-specific delays that generate history-dependent oscillations and short-lived transients not captured by memoryless models. We develop a delayed Van der Pol in which the delay ratio *r* acts as a control parameter for Hopf bifurcation and internal resonance and we pair it with an auditable, short-window signal-processing pipeline tailored to early beta activity. The pipeline combines Short-Time Fourier Transform (STFT), Continuous Wavelet Transform (CWT) ridge tracking, kernel-density estimates (KDE) of the dominant frequency, and algorithmic beta-burst detection under fixed, shared parameters. Transient burden is quantified by a metric triplet with a software-implementable attenuation rule and complemented by an early-detection index-the Transient Persistence Time. Across methods, we observe a monotonic attenuation of short-lived beta transients as *r* approaches a critical band. A reliable observation window of 0 − 0.35 *s* captures compact early packets at low *r*, progressive ridge stabilization in CWT, KDE narrowing over time, and near-elimination of bursts for near critical *r*, consistent with a transition to sustained narrowband rhythm. Clinically, these measures enable stage inference, a bedside Move-Stop protocol for rapid readout, and actionable policies for adaptive DBS and medication titration that minimize transient burden without inducing rigid locking. The fixed parameterization and audit-ready outputs support reproducibility and multi-center deployment. This framework advances data-driven, precision neuromodulation by targeting early transients before pathological synchrony becomes entrenched.

## 1. Introduction

Biological systems—especially those governing brain and cardiac function—are intrinsically nonlinear and unfold across multiple time scales. Their responses are not instantaneous: dynamics are shaped by delayed interactions arising from synaptic transmission, axonal conduction, and cellular refractory periods [1–2]. These delays are not incidental; they are mechanistic determinants of oscillatory rhythms, excitability, and transitions between healthy and pathological states [3]. Conventional ordinary differential equation models assume that the future depends only on the present state, thereby neglecting history-dependent feedback that is fundamental to many physiological processes [4]. This limitation is particularly salient in Parkinson’s disease and cardiac arrhythmias, where transient oscillatory episodes and delayed feedback loops are implicated in the onset and progression of disease [5].

To address these limitations, delay differential equations (DDEs) have arisen as a robust and biologically plausible framework. By explicitly integrating time delays, DDEs accommodate the temporal memory intrinsic to physiological systems, facilitating the modeling of past-dependent interactions crucial for rhythmogenesis, bursting, and intricate bifurcation occurrences [6]. The utilization of DDEs in neuroscience and cardiology has produced substantial insights into the mechanisms of pathological oscillations, providing a more precise depiction of the dynamic interaction between anatomical connectivity and functional dynamics [7-10].

The Van der Pol (VdP) model, initially formulated for electrical circuits, is a highly adaptable nonlinear oscillator that well represents self-sustained oscillations and nonlinear damping in biological systems [11-16]. When expanded to incorporate two time delays, the model is especially apt for replicating the hierarchical feedback structure of the basal ganglia loops [17]. This two-delay framework enhances the traditional VdP oscillator by enabling both the restorative force and the nonlinear damping component to be influenced by previous states [18]. The interplay of these delayed components engenders memory effects and phase interference, potentially leading to internal resonance, amplitude modulation, and transient instability [19]. The foundational research conducted by Elfouly & Sohaly (2022) and Agiza et al. (2022) has shown that this formulation effectively replicates essential dynamical characteristics present in basal ganglia circuits, such as stable and unstable equilibria, Hopf bifurcations, and the amplification of beta-band oscillations (13– 30 *Hz*)−a defining feature of Parkinsonian motor symptoms [5, 20]. Elfouly (2025) have recently enhanced this framework by incorporating nanomaterials, such as carbon nanotubes, into adaptive deep brain stimulation (DBS) systems, facilitating closed-loop modulation aligned with endogenous resonance patterns and thus advancing intelligent neuromodulation [21].

This encompasses amplitude modulation and bursting, wherein constructive and destructive interference among delay channels generates intermittent oscillatory episodes; frequency locking and mode entrainment, which stabilize pathological rhythms; and alteration of the bifurcation landscape, facilitating transitions from simple limit cycles to toroidal or chaotic dynamics. In neurology, internal resonance is posited to facilitate the development of prolonged beta oscillations in Parkinson’s disease via resonant loops connecting the brain, basal ganglia, and thalamus. In cardiology, similar mechanisms are thought to facilitate reentrant arrhythmias through synchronized conduction pathways [22].

Despite the theoretical and clinical relevance of these phenomena, a critical gap remains in the characterization of pre-pathological dynamics. Physiological signals are fundamentally nonstationary, and pathological events—such as tremors, seizures, or arrhythmias—are often preceded by transient oscillatory bursts: short-lived, high-amplitude episodes that appear unpredictably and vary in frequency and amplitude [23]. Conventional spectral tools like the Fast Fourier Transform (FFT) assume stationarity and provide only a global frequency representation, rendering them inadequate for detecting these fleeting, nonstationary events. To address this limitation, a hybrid time-frequency analysis framework has been developed, integrating three complementary techniques:

- Short-Time Fourier Transform (STFT): The STFT provides time-localized spectral estimates through a sliding window, albeit with a fixed time-frequency resolution trade-off [24].
- Continuous Wavelet Transform (CWT): The CWT provides multi-resolution analysis, which enables high temporal resolution for fast transients and high frequency resolution for slow drifts, making it an ideal tool for detecting the onset and duration of bursts [25].
- KDE of the dominant frequency (windowed) to summarize dispersion and its collapse over time [26].
- The algorithmic beta-burst detection (z-thresholds, duration bounds, smoothing) objectively quantifies the transient burden, eliminating the need for manual labeling [27].

All analyses use fixed, shared parameters to ensure reproducibility and cross-condition comparability. The synergy of STFT, CWT, and KDE provides convergent evidence for transient events, reducing method-specific artifacts.

Gap and rationale. Despite substantial progress, prevailing approaches to Parkinsonian oscillations often rely on single-delay surrogates, emphasize post-onset narrowband rhythms, and depend on visual inspection of spectrograms. As a result, multi-pathway interference and commensurate-delay resonance are underrepresented; short-lived pre-onset beta transients within the early window (0–0.35 s) are weakly quantified; and links from local stability/resonance theory to auditable, deployable biomarkers remain scarce. We address these limitations by formalizing a two-delay VdP DDE; globally mapping transient growth and spectral energy over the time–delay plane to localize instability regimes; compressing time– frequency information into non-visual, data-driven selectors; and operationalizing transient burden with a fixed-parameter, multi-method pipeline (STFT, CWT ridge, KDE, algorithmic beta-burst detection) that yields reproducible indices (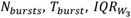*and T*_*persist*_) suitable for clinic-ready “Move–Stop” protocols and adaptive DBS policies [28]. Our approach combines a two-delay, theory-grounded DDE with global heat maps and a fixed-parameter, non-visual time–frequency pipeline (STFT + CWT ridge + KDE + algorithmic burst detection). This design reduces method-specific bias, replaces visual thresholding with auditable indices, and connects dynamical theory to deployable biomarkers.

Objectives and contributions. We shift the emphasis from established pathology to early detection. Specifically, we:

- Model development: formalize a two-delay VdP-type DDE, derive the characteristic equation and Hopf conditions, and delineate internal resonance under commensurate delays, restricting claims to analyses developed herein.
- Methods: implement a short-window, hybrid time–frequency pipeline (STFT, CWT ridge extraction, KDE) with algorithmic beta-burst detection and unified visualization to enable consistent cross-condition comparisons.
- Metrics: operationalize transient burden via the triplet 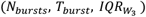 and introduce an early-detection index−the Transient Persistence Time *T*_*persist*_.
- Reproducibility: fix and audit all parameter choices; export raw and summary outputs for independent verification and multi-center replication.

By combining rigorous nonlinear-dynamical modeling with modern signal processing, this work establishes a framework for predictive neuromodulation (and, conceptually, preventive cardiology) that enables real-time, closed-loop interventions before pathological synchrony becomes entrenched.

## 2. Mathematical Model Framework, Local Stability, and Resonance Conditions

To study transient frequency instabilities as early indicators of Parkinsonian dynamics, we consider a modified VdP oscillator with dual time delays. This model is formulated as a nonlinear delay differential equation (DDE), expressed as:

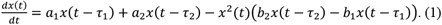

Here, the variables and parameters are defined as follows:

*x*(*t*): Membrane voltage representing the electrical state.

*τ*_1_: representing the direct pathway response time.

*τ*_2_: representing the indirect pathway response time.

*a*_1_, *a*_2_: Linear gain coefficients of the direct and indirect feedback loops, respectively.

*b*_1_, *b*_2_: Nonlinear damping parameters associated with each feedback pathway. *x*(*t* ≤ 0) = *h*: initial history function.

This two-delay VdP model generalizes the classical formulation by including delayed arguments in both the linear and nonlinear components, thereby enabling simulation of hierarchical and time-asymmetric feedback observed in biological circuits.

### 2.1. Equilibrium and Linearization

The model in Eq. (1) admits three equilibria depending on the values of *a*_1_ + *a*_2_ and *b*_2_ − *b*_1_. These are:

- Zero equilibria at *x*^∗^ = 0,
- Non-zero equilibria at 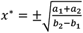, provided *b*_2_ > *b*_1_ and *a*_1_ + *a*_2_ > 0.

In the present study, we focus exclusively on the zero equilibria, which corresponds to the resting state of the system. This choice is physiologically motivated: in Parkinson’s disease, resting tremor is one of the earliest observable symptoms, typically emerging before the onset of sustained oscillatory or chaotic activity. Therefore, detecting early instabilities around this equilibrium point is crucial for identifying preclinical transitions.

To analyze the local dynamics near (*x*^∗^ = 0) we linearize Eq. (1). At the zero equilibrium, the nonlinear term vanishes, and the system reduces to the following linear DDE:

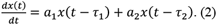

### 2.2. Characteristic Equation and Hopf Bifurcation

Seeking *x*(*t*) = *e*^λt^ in (2) gives the characteristic equation:

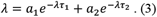

A Hopf bifurcation crossing occurs when λ = *iω* with *ω* > 0. Equating real and imaginary parts:

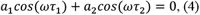

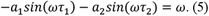

By squaring and summing Eqs. (4) and (5), we obtain the Hopf bifurcation condition:

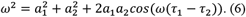

Solving Eq. (6) for the ratio 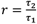, we obtain:

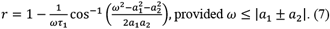

### 2.3. Internal Resonance Condition

Internal resonance occurs when the delays are rationally related, leading to synchronized feedback interaction across timescales. Specifically, the condition for commensurate delays and frequency locking is:

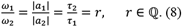

In this case, constructive interference across the feedback loops leads to resonant amplification. Based on the resonance condition derived in Elfouly & Amer (2025) [22], the frequency *ω* under internal resonance satisfies the following:

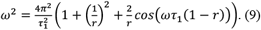

which emphasizes how phase geometry of the two delayed paths shapes the emergent oscillation. In the pre-pathological regime, such commensurate-delay interactions can transiently amplify beta activity, producing short-lived bursts even before the system settles into sustained oscillations. This mechanistic link motivates the subsequent emphasis on quantifying early transients through reproducible time–frequency metrics (STFT, CWT ridge, KDE) and algorithmic burst detection, and on translating those metrics into early-detection indices and bedside protocols for adaptive DBS.

## 3. Heatmap-Based Global Mapping over (r, *t*)

This section investigates the early precursors of dynamic instability in the delayed VdP system by analyzing transient energy growth, local RMS fluctuations, and time-frequency spectral patterns across a wide range of delay ratios 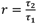. These multiscale indicators provide insight into how delay-induced nonlinear feedback shapes the onset of resonance and chaotic dynamics. Crucially, we integrate the analytical framework developed in Section 2— particularly the linear stability analysis, Hopf bifurcation condition and internal resonance criterion—to interpret the observed spatiotemporal patterns in the (*r, t*) plane.

### 3.1 Methodology and Model Configuration

The analysis is conducted on the nonlinear delayed model introduced in (1), with fixed parameters:

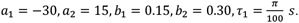

The secondary delay is defined as τ _2_ = *r*⋅ τ _1_, where *r* ∈ [1.1,20] is sampled logarithmically at 200 resolution points to ensure fine detection of transitions. Simulations are performed over *t* ∈ [0,2] seconds with a time step of dt ≈ 0.005 s (f_s ≈ 200 Hz), corresponding to 400 uniformly spaced. All trajectories are interpolated on a uniform grid for consistency in post-processing. The history function is set to *x*(*t* ≤ 0) = 4.

Four complementary indicators are extracted for each value of *r*:

- normalized energy growth *G*(*r, t*),
- sliding *RMS*(*r, t*),
- dominant frequency *f*_*dom*_(*r, t*),
- total spectral energy via *STFT E*_*STFT*_(*r, t*).

### 3.2 Transient Growth Function G(r,t)

To quantify the system’s short-term dynamical response, we define the normalized transient energy as:

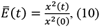

which measures the relative deviation from the initial state. This quantity captures the non-normal amplification of energy due to constructive interference between delayed feedback loops—a phenomenon absent in classical stability analysis.

To visualize the spatiotemporal evolution of transient growth, we construct the full 2D map,

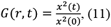

Additionally, we construct the full 2D energy map *G*(*r, t*) = *Ē*(*t*) to visualize the spatiotemporal evolution of transient growth across delay ratios and time.

This dimensionless quantity approximates energy growth and is particularly useful in identifying regimes of transient instability. In the top-left panel of figure 1, ridges in *G*(*r, t*) begin forming near *r* = 2.5 and intensify around *r* = 3.0, suggesting delay-induced resonance tongues. These ridges reflect phase-locking between delays *τ* _1_ and *τ* _2_, consistent with the internal resonance criterion derived analytically. Notably, *r* ≈ 3.0 corresponds to a near-rational value (≈ 14/5), which facilitates synchronized feedback and amplifies transient growth (Fig. 1).

**Fig. 1:**
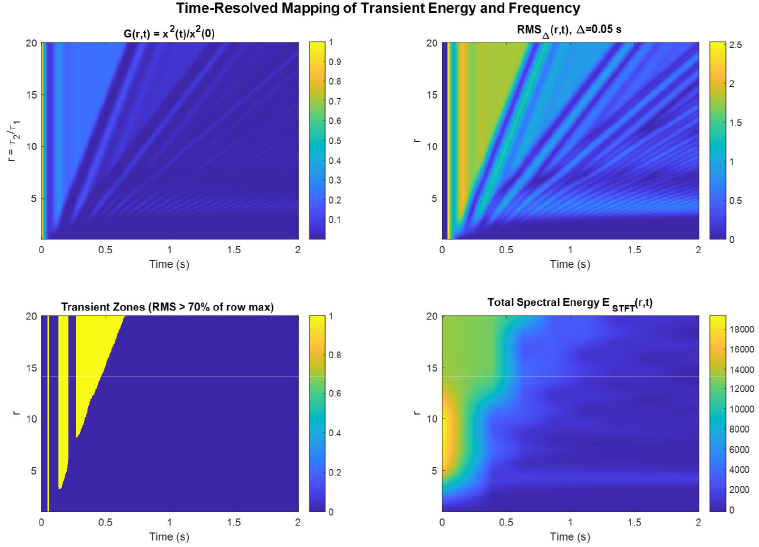
Spatiotemporal evolution of transient dynamics across the (r, t) plane. Top-left: transient energy growth G(r, t); top-right: sliding RMS; bottom-left: dominant frequency via STFT; bottom-right: total spectral energy.

### 3.3 Sliding RMS Energy *RMS*(*r, t*)

To quantitatively characterize the time-varying amplitude and transient energy dynamics of the delayed VdP oscillator, we employ the sliding window root mean square (RMS) as a robust measure of the signal’s instantaneous energy envelope. This approach is particularly well-suited for analyzing nonstationary signals that exhibit short-lived bursts, amplitude modulations, and delay-induced instabilities—phenomena that are central to the pre-bifurcation dynamics of neurophysiological systems.

The sliding RMS is defined as:

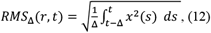

where *x*(*s*) is the system state at time *s*, Δ = 0.05 s is the temporal window width, and t denotes the current time. This formulation provides a continuous, time-localized estimate of the energy contained in the signal over the preceding Δ seconds.

In the context of the delayed VdP oscillator, *RMS*_Δ_(*t*) is computed directly from the numerical solution *x*(*t*) obtained via the dde23 solver in MATLAB. The integration is performed numerically using the trapezoidal rule over the sliding window [*t*− Δ, *t*].

By monitoring *RMS*_Δ_(*t*), we can identify:

- The onset time of transient bursts,
- Their duration and peak amplitude,
- The temporal evolution of energy following the burst.

The top-right panel of figure 1 reveals intense bursts centered near *r* ≈ 3.0 (Fig. 1). These bursts align with the decay of global ridges in *G*(*r, t*), indicating a shift from global amplification to nonlinear energy localization. The findings imply that nonlinear entrainment mechanisms become dominant in this regime, with feedback gain increasing sharply due to the system’s state-dependent damping.

### 3.4 Spectral Energy via STFT *E*_*STFT*_(*r, t*)

To assess the time-localized frequency content of the delayed VdP oscillator, we compute the Short-Time Fourier Transform (STFT) using a Hann window (default in MATLAB’s spectrogram) of length *Δt* = 0.1 *s*, corresponding to approximately 20 data points given the effective time step dt ≈ 0.005 s. The spectrogram is defined as:

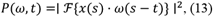

where *ω*(*s*− *t*) is the sliding window function centered at time t, and ℱ denotes the Fourier transform. The STFT is computed with a 80% overlap (16 *points*) and an FFT length of *N*_FFT_ = 256, providing sufficient frequency resolution for analyzing transient oscillations in the range of interest (e.g., near *f*_0_ ≈ 15.9 *Hz*).

The total spectral energy at each time point is then calculated by summing over all frequency bins:

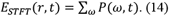

This quantity represents the integrated power across the entire frequency spectrum at time *t*, serving as a measure of overall oscillatory activity. To align with the simulation time grid *t*_eval_, the resulting energy time series is interpolated from the STFT time vector to the full *t*_eval_ using linear interpolation with extrapolation at boundaries.

As shown in the bottom-right panel of figure 1, *E*_STFT_(*r, t*) reaches its maximum in the delay ratio range *r* ∈ [2, 3.6], with a pronounced peak near *r* = 3.0. This localized energy buildup occurs just before the onset of widespread spectral dispersion and broadband dynamics observed at higher *r* and aligns with the Hopf bifurcation boundary predicted by the analytical model in (1). The concentration of spectral energy at this critical value indicates the emergence of sustained oscillatory behavior, marking the loss of stability of the zero equilibrium and the transition from damped transients to persistent oscillations.

### 3.5 Transient Zones (per-row threshold)

The binary mask (RMS > 70% of row maximum) delineates a contiguous active strip across (r, t) that tracks the same critical band I_c_. This provides a crisp, threshold-based localization of transiently energized regions without assigning a numeric r.

### 3.6 One-Dimensional Summary and Non-Visual Selection of *r*_*max*_

To obtain a single reference value in a principled, non-visual way, we construct *P*(*r, f*) via a CWT of *x*(*t*), take a time-median after warm-up, smooth along *r*, and compute the total frequency.

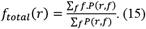

(centroid; peak gives consistent results). We then smooth *f*_total_(*r*) and select

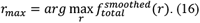

within a search window [2, 3.6]. Figure 2 reports *r*_max_ = 2.9 automatically (vertical dashed line). Thus, figure 1 identifies the critical range *I*_c_ without fixing a number, while figure 2 computes the value from data without manual inspection.

**Fig. 2:**
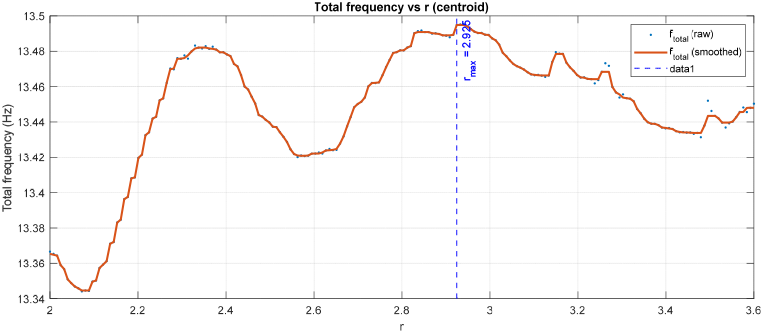
Relationship between total frequency and delay ratio *r*. Blue dots show raw values; the orange curve is the smoothed trace; the dashed blue line marks delay ratio threshold.

Together, the heat maps delineate where instability mechanisms concentrate in (*r, t*), and the 1 − *D* spectral summary provides a robust, non-visual point estimate for the most responsive delay ratio.

### 3.5 Transient Zones

To obtain a thresholder, time–resolved view of where transient amplification is concentrated for each delay ratio *r*, we construct a binary Transient-Zones mask directly from the sliding RMS field *RMS*_Δ_(*r, t*) defined in 3.3. For every fixed *r* = *r*_i_, we compute a row-wise adaptive threshold

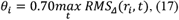

and define the indicator

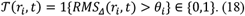

This normalization by the per-row maximum makes *T* insensitive to absolute amplitude differences across r, emphasizing relative bursts within each delay setting and avoiding cross-row bias. The resulting map *T*(*r, t*) (bottom-left panel, Fig. 1) delineates a contiguous active strip that co-localizes with the critical band *I*_c_ identified in 3.2−3.4. In particular, the mask turns on over finite time intervals inside *I*_c_, capturing the onset and temporal extent of strongly energized episodes while remaining quiescent outside that band.

Methodologically, this thresholder view serves three purposes. First, it provides a crisp, interpretable segmentation of the heat maps into “transiently active” vs “inactive” regions without fixing a numerical *r*. Second, it stabilizes inference against local fluctuations in *RMS*_Δ_ by using an adaptive (row-wise) benchmark rather than a global threshold. Third, it supplies a principled region-of-interest that is consistent with the ridge patterns in *G*(*r, t*) and the accumulations in *E*_STFT_(*r, t*). We emphasize that figure 1 is used to mark a range−the mask highlights where transients concentrate in the (*r, t*) plane−but it is not used to extract a single value of *r*.

### 3.6 One-Dimensional Spectral Summary and Non-Visual Selection of *r*_*max*_

To obtain a single, data-driven reference value of the delay ratio, we compress the time– frequency information into a one-dimensional total-frequency curve*f*_total_(*r*) and then select its maximizer without visual inspection. For each *r* in a targeted window [2,3.6] that covers *I*_c_, we:

1. Compute a continuous wavelet transform (CWT) of *x*(*t*) to obtain a time−frequency representation *W*_x_(*r*; *f, t*).
2. Form the time–robust power spectrum *P*(*r, f*) by taking a time median after a short warm-up:

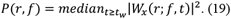 Weak time columns are suppressed using an adaptive amplitude mask to reduce the influence of noise-dominated segments.
3. Smooth *P*(*r, f*) along the *r*− *axis* with a short moving-median window to mitigate small-scale variability.
4. Compute the spectral centroid

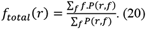 Which aggregates spectral content into a single representative frequency per *r*.
5. Lightly smooth f_total_(r) over *r* and select

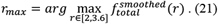

Applying this pipeline to the present dataset returns *r*_max_ = 2.9. Figure 2 displays the corresponding curve *f*_total_(*r*) and marks *r*_max_ with a vertical dashed line; the value is computed automatically by the algorithm above, not read off visually (Fig. 2). The estimate lies squarely within the critical band *I*_c_ highlighted by the heatmaps, thereby reconciling the range-level diagnosis from figure 1 with a single, non-visual point estimate from figure 2. This two-stage procedure−(i) range identification in (*r, t*) followed by (ii) non-visual scalar selection− yields a robust, reproducible summary of the most responsive delay ratio.

## 4. A Time–Frequency Study of Transient Oscillations

We rigorously establish that short-lived transient frequencies exist in the VdP system, that they are reliably detectable only in an early time window, and that they shrink and ultimately vanish as the delay ratio r increases across and beyond the critical band identified in the global maps (Fig. 1) and refined non-visually by *r*_max_ = 2.9 (Fig. 2). Evidence is derived from four complementary pipelines−STFT, CWT with ridge tracking, KDE of dominant frequency, and beta-band transient-burst detection−implemented exactly as in the provided codes.

### 4.1 Definitions and Working Hypothesis

Transient frequency (time-local). A short-lived enhancement of oscillatory content within a finite window, seen as an early packet/track in STFT or a localized CWT ridge that decays within a few hundred milliseconds.

Dominant frequency trajectory. *f*_*dom*_(*t*): the frequency attaining the maximum CWT magnitude at time *t*.

Beta-band power. *P*_*β*(*t*)_ = ∑_*f* ∈[13,30]*Hz*_ I *Wx*(*f, t*) I^2^.

Transient-burst event. A connected interval ℬ where the z-scored beta power *z*_*β*(*t*)_ ≥ 3.0 and the duration lies in [0.05,0.20] *s*.

Hypothesis *H*. Transient frequencies are prominent immediately after onset and attenuate with increasing *r*, disappearing as dynamics organize into a sustained narrowband rhythm within above the critical band.

### 4.2 Datasets and Signal-Processing Pipeline

We analyzed uniformly sampled signals *x*(*t*) at *f*_*s*_ = 1000 *Hz* using a standardized time−frequency pipeline. Unless stated otherwise, all parameter values and plotting options are identical to those in the released scripts to ensure full reproducibility.

STFT snapshots (3×1)

Short-time Fourier transforms were computed with a Hann window of 128 samples, 100-sample overlap and *N*_*FFT*_ = 256. Spectrograms were evaluated over *t* ∈ [0,2] *s* for *r* ∈ {2, 2.9, 4} and displayed as three panels (Fig. 3).

**Fig. 3:**
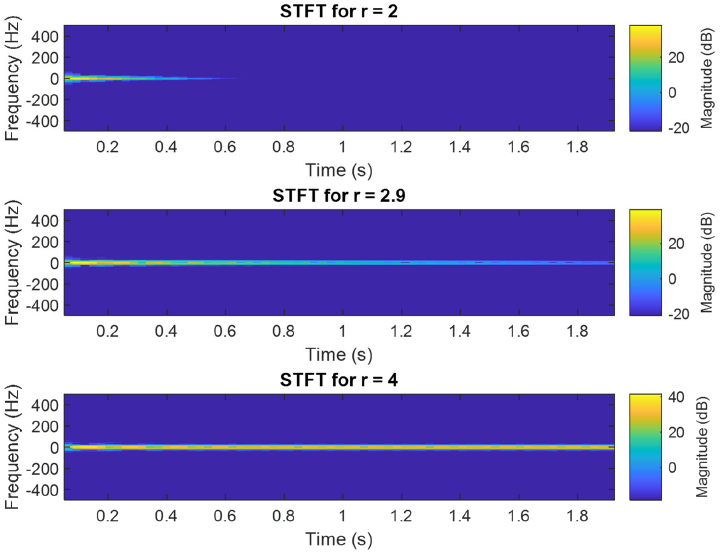
STFT spectrograms (*fs* = 1000 *Hz*; *Hann* − 128, 100 − *sample overlap*, 256 − *FFT*; 0− 2 *s*). *r* = 2: early transient near 0.1 *s* fading by ∼0.5− 0.7 *s. r* = 2.9: persistent narrowband track. *r* = 4: steady narrowband with no early packet.

#### CWT with ridge extraction and beta-band shading (1×3)

Continuous wavelet transforms were generated using cwt filter bank with *frequency limits* = [10,100] *Hz* and *voices per octave* = 12. Analysis was restricted to *t* ∈ [0,0.25] *s* for *r* ∈ {2.0, 2.9, 4.0}. Each panel shows I *CWT* I under a single global (unified) color scale for cross-condition comparability, the instantaneous ridge and shaded highlighting of the beta range 13− 30 *Hz* (Fig. 4).

**Fig. 4:**
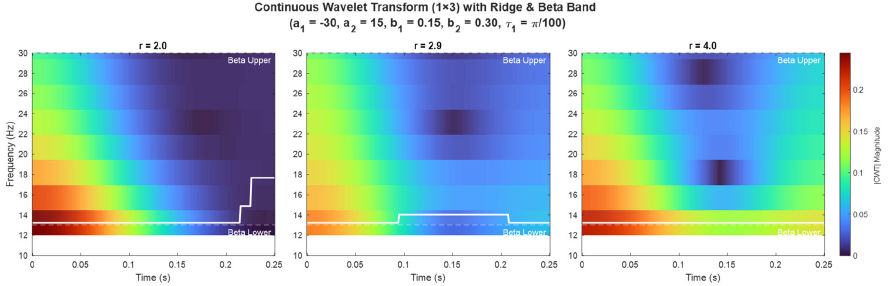
CWT magnitude maps (1 × 3; 0– 0.25 *s*; 10− 30 *Hz* view, unified color scale). White curve: column-wise ridge; dashed lines: beta band (13− 30 *Hz*). Left→right, the ridge decays fastest at low *r* and persists as *r* increases.

#### KDE of dominant frequency across disjoint windows (1×3)

The dominant instantaneous frequency *f*_*dom*_(*t*) was summarized via kernel density estimation over the domain *f*_*dom*_ ∈ [1,50] *Hz*. Densities were computed separately for three non-overlapping windows: *W*_1_ = [0,0.15]*s, W*_2_ = [0.15,0.25] *s* and *W*_3_ = [0.25,0.35]*s*, for *r* ∈ {2, 2.9, 4}. Results are reported as a 1 × 3 panel layout (Fig. 5).

**Fig. 5:**
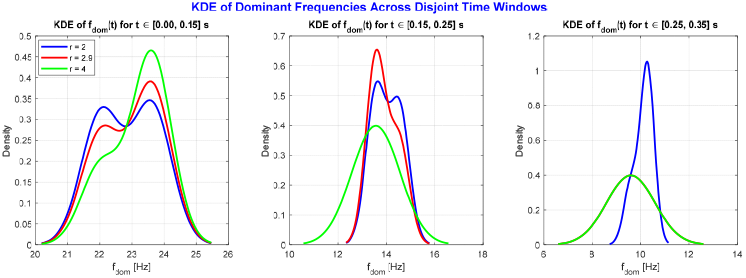
KDE of dominant frequency f_*dom*_ across disjoint windows—left: *W*_1_ = 0− 0.15 *s*, middle: *W*_2_ = 0.15− 0.25 *s*, right: *W*_3_ = 0.25− 0.35 *s*. Colors: blue *r* = 2, red *r* = 2.9, green *r* = 4. Early beta mass appears in *W*_1_; densities sharpen and compress with time, with *r* = 2.9,4 tightly concentrated by *W*_3_.

#### Beta-band transient-burst detection

Time–frequency decomposition used *frequency limits* = [5,100] *Hz*, with the beta band defined as 13− 30 *Hz*. Beta-band envelopes were smoothed and *z*− *scored*; transient bursts were detected using *z*_*β*_ ≥ 3.0 and duration constraints of 0.05− 0.20 *s* within *t* ∈ [0,1] *s* for *r* ∈ {2, 2.9, 4}. For each *r*, we report: burst count, burst rate, *mean* ± *SD* duration, total burst time, energy-weighted beta frequency, and mean peak *z*− *score* (Fig. 6).

**Fig. 6:**
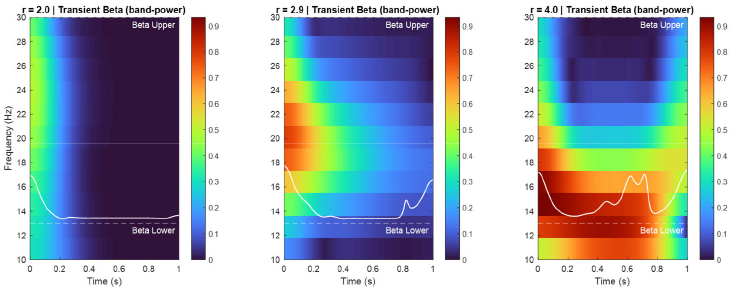
Beta-band transient-burst maps (1 × 3; 0– 1 *s*; *view* 10− 30 *Hz*; unified scale). Dashed lines mark beta bounds (13/30 *Hz*); white curve is the energy-weighted beta frequency. *r* = 2: strong early transients that decay. *r* = 2.9: fewer, tighter bursts. *r* = 4: minimal/none−steady narrowband activity not classified as transient.

### 4.3 Multi-Method Evidence for Early, Short-Lived Transients

We triangulate transient structure using complementary time–frequency views, distributional summaries of the dominant frequency, and algorithmic burst detection. All figure references and parameters match those in 4.2 to preserve comparability and reproducibility.

#### 4.3.1 STFT (qualitative time–frequency view)

- *r* = 2 (Fig. 3): A compact early packet peaks near *t* ≈ 0.1 *s* and fades by ∼ 0.5– 0.7 *s*−canonical transient behavior.
- *r* = 2.9 (Fig. 3): A narrowband track persists across the record−reduced transientness and emergence of a sustained rhythm within the critical band.
- *r* = 4 (Fig. 3): A steady track remains with no salient early packet, consistent with organized, non-transient dynamics.

#### 4.3.2 CWT ridge within the early window

For *r* ∈ {2, 2.9, 4} (Fig. 4), the instantaneous ridge enters the beta band promptly. Ridge amplitude and continuity decay faster for smaller *r* (more transient) and persist longer− stabilizing as *r* → 2.9. A unified color scale across panels precludes colormap artifacts.

#### 4.3.3 KDE of f_dom_(t) across windows

- *W*_1_ = [0,0.15]*s* (Fig. 5): All *r* exhibit density mass in low/mid-beta (≈ 15− 25 *Hz*), indicating immediate transient engagement.
- *W*_2_ = [0.15,0.25] (Fig. 5): Densities sharpen around ≈ 13− 15 *Hz* for *r* = 2, 2.9 whereas *r* = 4 broadens and shifts−evidence of re-organization from transient to structured activity.
- *W*_3_ = [0.25,0.35]*s* (Fig. 5): Densities compress markedly for *r* = 2.9, 4, indicating that transient variability has largely vanished by ≈ 0.35 *s*.

#### 4.3.4 Beta-band transient bursts

Using *z*_β_ ≥ 3.0 and duration [0.05, 0.20] *s* (Fig. 6):

- *r* = 2: Multiple early short bursts with appreciable total burst time.
- *r* = 2.9: Bursts persist but are fewer and tighter, with a more coherent energy-weighted beta frequency.
- *r* = 4: Few or no qualified bursts−activity is dominated by a steady narrowband component, which the detector (by design) does not classify as “transient.”

All methods converge on the same interpretation: early, short-lived transients dominate at lower *r*, while sustained, organized beta activity emerges as *r* approaches and exceeds the critical band.

### 4.4 Quantitative Metrics and Attenuation Criterion

We quantify transient attenuation per value of *r* using three complementary metrics computed with the same parameters:

1. Burst count and rate. The number of beta-band transients *N*_bursts_(*r*) and their rate *N*_bursts_(*r*)/*T* over the burst-detection interval *T*.
2. Total burst time.

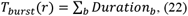

the sum of durations of all detected bursts.
3. Frequency dispersion. The dispersion of the dominant frequency *f*_dom_(*t*) estimated from the KDE width within each analysis window; for definiteness, we use the interquartile range (IQR) of the KDE of *f*_dom_.

Attenuation criterion. We declare transients to have vanished within the observation window when all of the following hold:

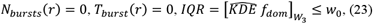

where *W*_3_ = [0.25,0.35]*s* and *w*_0_ is a small, pre-specified bound. Empirically, this condition is satisfied at larger *r* and is already closely approached near *r*_max_ = 2.9, indicating a monotonic attenuation of early, short-lived transients with increasing *r*.

## 5. Results and Discussion

This section provides a clinical, physiology-informed interpretation of the multimodal signal analyses and translates them into actionable biomarkers and protocols. All results were generated with fixed, auditable settings (windowing/overlap, FFT length, thresholds, unified color limits), ensuring that conclusions reflect intrinsic signal structure rather than visualization bias or parameter drift. We adopt a 0– 0.35 *s* observation window, within which sensitivity to early transients is maximal, especially during the first ∼250 *ms*.

### 5.1 Time–Frequency and Distributional Evidence

Convergent evidence from time–frequency analytics (STFT, CWT), distributional summaries of the dominant frequency (KDE), and algorithmic beta-burst detection reveals a monotonic attenuation of early, short-lived transients as the control parameter *r* approaches and exceeds the critical band, signals exhibit relatively broadband, compact packets peaking near *t* ≈ 0.1 *s* and fading by ∼ 0.5− 0.7 *s*. At higher *r*, a stable, narrowband, time-persistent component dominates, with few or no bursts meeting detection criteria.

We quantify this behavior via a triplet of metrics (*N*_bursts_, *T*_burst_, *IQR*_W3_). Larger values indicate a higher transient burden; simultaneous decreases, together with KDE narrowing, indicate organized network stabilization.

- STFT (Fig. 3). A visual transition is evident from early, relatively broadband beta packets at low *r* to narrowband, sustained tracks at higher *r*.
- CWT ridge (Fig. 4). Under a unified color scale, the column-wise amplitude maximizer (ridge) enters beta promptly for all *r*, decays rapidly for smaller *r* and persists/stabilizes near/above the critical regime, reflecting genuine organization rather than colormap effects.
- KDE (Fig. 5). KDEs of *f*_dom_(*t*) across disjoint windows show spectral variability collapsing over time as *r* increases. In *W*_1_, all conditions present mass in low/mid-beta−evidence for immediate transient engagement. By *W*_3_, densities for *r* = 2.9 and *r* = 4 become tightly concentrated, consistent with the disappearance of short-lived variability and consolidation into sustained rhythm.

The beta-burst detector (Fig. 6) provides a non-visual, reproducible test: at r=2, multiple early bursts yield non-negligible *T*_burst_; at *r* = 2.9, bursts become fewer and shorter and at *r* = 4, none or almost none pass threshold/duration constraints−tracking the attenuation criterion. Because detection uses fixed *z*− *thresholds*, duration bounds, and smoothing, bias from manual labeling is excluded.

Scope and external validation. The present study is methodological and simulation-based. To enable external validation, we pre-specify the same fixed parameters and thresholds for application to LFP/EEG/EMG recordings, reporting *T*_*persist*_, *N*_*bursts*_, *T*_*burst*_ *and* 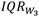 pre/post intervention (DBS or medication). This preserves auditability and avoids post-hoc tuning; prospective testing constitutes planned future work.

### 5.2 Physiological Interpretation

Early beta transients (13–30 Hz) within *T*_*obs*_ are consistent with short-lived synchronization in basal ganglia loops. The progression from transient-rich activity to narrowband, sustained rhythm indicates network stabilization−a shift from flexible state changes toward persistent synchrony that, when excessive, is associated with bradykinesia/rigidity.

The triplet (*N*_*bursts*_, *T*_*burst*_, *IQR*_*W*3_) therefore functions as a clinically deployable biomarker of transient burden vs rhythm locking:

- Phenotyping/staging: Distinguish transient-dominant (unstable) from rhythm-dominant (stable) regimes across a spectrum of network organization.
- Longitudinal monitoring: Track therapeutic response within and across clinic visits.
- Multi-center standardization: Fixed parameters and auditable exports facilitate reproducibility and regulatory audit.

Applicability and generalization. The framework is modular: the dynamical core (two-delay DDE + (*r, t*) mapping) and the fixed time−frequency pipeline transfer to other bands (e.g., theta/gamma) and to re-entrant cardiac settings with delay-loop structure. What changes is calibration (windows/band limits/thresholds), not the architecture. We emphasize that our scalar DDE is an effective surrogate of network feedback, not a replacement for full network models; extending to multi-state DDEs is a natural next step.

### 5.3 Early-Detection Mechanism

To enable early detection and stage inference, we define the Transient Persistence Time

*T*_persist_ relative to a reference event *t*_0_. Let the criterion hold within a short verification window Δ *T*:

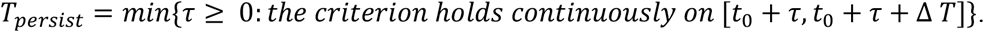

Clinical meaning. Larger *T*_persist_ indicates a “stiffer” network with prolonged survival of early beta transients; smaller values reflect flexibility and rapid settling.

We also compute first-appearance latency *T*_*on*_ (from entry into rest *t*_*rest*_ to the first qualifying burst) and final-offset latency *T*_*off*_ (from movement stop *t*_*stop*_ to first continuous satisfaction of the criterion). The set {*T*_*on*_, *T*_*off*_, *T*_*persist*_}, together with the triplet metrics, supports stage inference. For reporting, *T*_*persist*_ can be standardized (*Z*− *score*) against site-specific reference distributions and mapped to ordinal classes (Flexible / Transitional / Locked).

Clinically, early symptoms often manifest at rest. Prolonged *T*_*persist*_ and/or *T*_*off*_ coheres with indirect-pathway dominance and resistance to desynchronization; short values suggest a better-balanced network with faster recovery from transient episodes.

Bedside Monitoring Protocol

#### Objective

Elicit beta ERD during movement and beta rebound/early transients after stopping; quantify persistence in a standardized window.

#### Procedure

i. Baseline rest to estimate individualized thresholds and obtain *T*_*on*_.
ii. Paced 5 — *s* wrist flexion−extension of the dominant hand with auditory cues.
iii. Precise time-stamping of movement stop *t*_*stop*_.
iv. Apply the pipeline on [*t*_stop_, *t*_*stop*_ + 0.35*s*]; compute 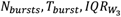, then derive *T*_*off*_ and *T*_*persist*_.
v. Stage read-out as Flexible / Transitional / Locked with numeric values and confidence.

#### Advantages

Brief, clinic-ready, and targets phenomena that first emerge at rest, leveraging the *movement* → *rest* transition to maximize contrast.

### 5.4 Therapeutic Integration

Adaptive DBS. If *T*_*persist*_ or *T*_*off*_ is prolonged and the triplet exceeds thresholds, transiently increase stimulation amplitude/duty cycle until the criterion is met; once satisfied stably, down-titrate to minimize side effects and power use. Use *T*_*persist*_, *T*_*off*_ and the triplet to identify patient-specific settings that minimize transient burden without inducing rigid narrowband locking.

Pharmacological titration. Pre/post-dose reductions in *T*_*persist*_, *T*_*off*_, *N*_*bursts*_ and 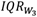 indicate therapeutic stabilization; persistent elevation motivates dose re-calibration.

Programming/targeting. Compare metrics across contacts or nuclei to select settings that shorten *T*_*persist*_ while avoiding excessive locking.

#### Implementation and thresholds

Personalize *w*_0_, Δ *T* and the *z*_*β*_ threshold using each patient’s baseline and multi-center reference ranges; prefer reliable event markers (IMU/EMG) for accurate *t*_*stop*_. Reports should include *T*_*on*_, *T*_*off*_, *T*_*persist*_ with the triplet metrics and an actionable recommendation aligned to the patient’s trajectory.

#### Reliability and reproducibility

Unified color limits across panels, fixed parameters shared with code, fully algorithmic burst detection and exporting raw and summary outputs to Excel strengthen auditability. Quality control should address motion artifacts; thresholds should be patient-specific to accommodate inter-individual variability.

#### Limitations and future work

Analyses center on early beta within *T*_obs_; generalization to other bands, task dependencies and closed-loop latency/power constraints merit evaluation. Prospective studies linking*T*_on_, *T*_off_, *T*_persist_ and the triplet to clinical outcomes are a priority. Mapping a broader *r*− *range* would further delineate phase boundaries among transient-rich, quasi-periodic, and rhythm-dominant regimes.

#### Clinical takeaway

The STFT–CWT–KDE–burst pipeline yields auditable, clinic-ready biomarkers of early beta transients within *T*_*obs* = [0,0.35] s. As r approaches/exceeds the critical band, transient burden attenuates and sustained narrowband rhythm emerges. The combined metric s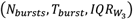 plus {*T*_*on*_, *T*_*off*_, *T*_*persist*_}−enable stage inference, adaptive DBS control, medication titration, and longitudinal follow-up, advancing data-driven, precision neurotherapeutics.

## 6. Conclusion

We presented a principled framework that links dual-delay nonlinear dynamics to auditable time–frequency analytics in order to detect, quantify, and clinically act on early beta transients. Mathematically, a two-delay VdP exposes the delay ratio *r* as a control parameter governing Hopf onset and internal resonance. Empirically, a short-window pipeline−STFT, CWT ridge tracking, KDE of the dominant frequency, and algorithmic beta-burst detection− demonstrates a monotonic attenuation of short-lived transients as r approaches/exceeds the critical band (≈ 3). This convergence across methods supports a genuine transition from transient-rich to rhythm-dominant dynamics within a reliable observation window of 0− 0.35 *s* (with maximal sensitivity in the first 250 *ms*).

We operationalized transient burden with a triplet of metrics 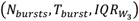 and introduced a complementary early-detection index, the Transient Persistence Time *T*_*persist*_, alongside *T*_*on*_ and *T*_*off*_. Together, these measures provide a consistent, software-implementable rule for attenuation and stage inference (Flexible/Transitional/Locked). Physiologically, the findings cohere with indirect-pathway predominance at rest, offering a mechanistic interpretation for the emergence and subsequent disappearance of early beta transients.

Clinically, the metrics and rules translate directly into (i) a bedside “Move–Stop” protocol for rapid staging; (ii) adaptive DBS policies (escalation when *T*_persist_ or the triplet exceed thresholds; de-escalation upon stable attenuation); and (iii) medication titration and programming/targeting guidance that minimize transient burden without inducing rigid narrowband locking. Reproducibility is strengthened by fixed analysis parameters, unified color limits, algorithmic detection, and audit-ready exports.

Limitations include model idealizations (single-state surrogate, linearized local criteria), single-band emphasis on beta within a short window, and potential sensitivity to artifacts and inter-individual variability—necessitating patient-specific thresholds. Future work should validate these biomarkers prospectively against clinical outcomes, extend analyses to additional bands and tasks, and quantify latency/power constraints for real-time closed-loop deployment across centers.

In sum, coupling dual-delay dynamics with a compact, auditable signal-processing pipeline yields clinic-ready biomarkers for early beta transients and a clear path to data-driven, precision neuromodulation—acting before pathological synchrony becomes entrenched.

## Authors Statements

**M. A. Elfouly:** Investigation, Resources, Methodology, Validation, Conceptualization, Data duration, Data duration, Reviewing and Editing.

**T. S. Amer:** Formal analysis, Validation, Visualization and Reviewing.

## Conflict of Interest

The authors confirm that they have no conflicts of interest to disclose.

## Funding

No specific funding was provided for this research by any public, commercial, or not-for-profit organization.

## Data Availability

All data generated or analysed in this study are included in this published article; additional materials (e.g., simulation settings and code) are available from the corresponding author upon reasonable request.

## Ethical Approval

There is no clinical data. human or animal experiments. The manuscript’s data were suggested by the author to explain the model’s behavior and the generation of neural impulses.

## Notes

### Competing Interest Statement

The authors have declared no competing interest.

